# SUMO Activated Target Traps (SATTs) enable the identification of a comprehensive E3-specific SUMO proteome

**DOI:** 10.1101/2022.06.22.497173

**Authors:** Daniel Salas-Lloret, Coen van der Meulen, Easa Nagamalleswari, Ekaterina Gracheva, Arnoud H. de Ru, H. Anne Marie Otte, Peter A. van Veelen, Andrea Pichler, Joachim Goedhart, Alfred C.O. Vertegaal, Román González-Prieto

## Abstract

Ubiquitin and ubiquitin-like conjugation cascades consist of dedicated E1, E2 and E3 enzymes with E3s providing substrate specificity. Mass spectrometry-based approaches have enabled the identification of more than 60,000 acceptor sites for ubiquitin and 40,000 acceptor sites for SUMO2/3. However, E3-to-target wiring is poorly understood. The limited number of SUMO E3s provides the unique opportunity to systematically study E3-substrate wiring. We developed SUMO Activated Target Traps (SATTs) and systematically identified substrates for eight different SUMO E3s, PIAS1, PIAS2, PIAS3, PIAS4, NSMCE2, ZNF451, LAZSUL(ZNF451-3) and ZMIZ2. SATTs enabled us to identify 590 SUMO1 and 1195 SUMO2/3 targets in an E3-specific manner. We found pronounced E3 substrate preference, even at the substrate isoform level. Quantitative proteomics enabled us to measure substrate specificity of E3s, quantified using the SATT index. Furthermore, we developed the Polar SATTs web-based tool (https://amsterdamstudygroup.shinyapps.io/PolaRVolcaNoseR/) to browse the dataset in an interactive manner, increasing the accessibility of this resource for the community. Overall, we uncover E3-to-target wiring of 1681 SUMO substrates, highlighting unique and overlapping sets of substrates for eight different SUMO E3 ligases.

## INTRODUCTION

Protein fate and function is controlled by numerous Post-Translational Modifications (PTMs). Among them, ubiquitination is the second most abundant PTM after phosphorylation, and controls virtually every process in eukaryotic cells in a dynamic manner. Ubiquitination consists of the covalent attachment of the small 76 amino acids ubiquitin protein to acceptor proteins and is performed by an enzymatic cascade in which ubiquitin-activating enzymes (E1) activate ubiquitin and transfer it to ubiquitin-conjugating enzyme (E2) which conjugates ubiquitin to the substrate assisted by an ubiquitin-ligase enzyme (E3). E3s are responsible for determining substrate specificity. The human genome encodes for two ubiquitin E1s, 30-40 E2s and more than 600 E3s^1^.

Similar to ubiquitin, other ubiquitin-like (Ubl) modifiers exist, which have dedicated E1-E2-E3 enzymatic cascades. Among these Ubls, Small Ubiquitin-like Modifiers (SUMOs) are the most abundant ones after ubiquitin. In vertebrates, there are three different types of active SUMOs, SUMO1, SUMO2 and SUMO3. Mature SUMO2 and SUMO3 differ only in a couple of amino acids and are commonly referred to as SUMO2/3. In contrast to ubiquitin, vertebrates express a single E1, a single E2 and less than a dozen *bona fide* E3s for SUMOs^1^.

Recent advances in mass spectrometry technologies and the optimization of sample preparation methodologies^2^ have enabled the identification of several tens of thousands of acceptor sites on thousands of proteins in human cells both for ubiquitin and SUMOs ^3–9^. However, our knowledge on E3-substrate wiring is still very limited. Determining which E3 modifies which substrate is a major challenge.

For ubiquitin, given the high number of E3s, solving the E3-to-target wiring in a proteome-wide manner is virtually impossible. However, for SUMOs, the E3 complexity is limited, simplifying this task. A proposed approach has been the quantification of changes on the SUMO proteome after SUMO E3 overexpression^10^, which, in principle, is an indirect measure. Another applied approach has been the performance of sumoylation assays on protein array-based screens ^11^, which is an *ex vivo* system that misses out on the restricted subcellular localization of proteins and lacks protein-protein complexes that are abundant in cells.

Here, we took advantage of our previous experience in the systematic identification of ubiquitination substrates using Ubiquitin Activated Interaction Traps (UbAITs)^12^ in the Targets of Ubiquitin Ligase Identified by Proteomics (TULIP) methodologies ^13–15^ and applied it for the identification of SUMO E3-specific substrates in a systematic manner for SUMO E3s in a proteome-wide approach.

## RESULTS

### SUMO E3 overexpression causes SUMO2/3 depletion in an RNF4-dependent manner

Aiming to identify putative E3-specific sumoylation substrates, we employed a similar approach as previously done with PIAS1^10^. We transiently transfected GFP-tagged constructs for the SUMO E3s PIAS1, PIAS2, PIAS3, PIAS4, ZNF451 and the Lap2α isoform of the ZNF451 SUMO Ligase (LAZSUL) in U2OS cells. To evaluate the transfection efficiency of our constructs, we analyzed our cells by fluorescence microscopy after immunostaining for SUMO2/3 (Figure 1A, Supplementary Figure 1). GFP-positive cells could be observed for every construct at different efficiencies, except for GFP-PIAS2, which transfection did not lead to the appearance of GFP-positive cells. Unexpectedly, the immunofluorescence SUMO2/3 signal was highly reduced in GFP-positive cells for PIAS1, ZNF451 and LAZSUL (Figure 1A, Supplementary Figure 1). Therefore, we quantified the SUMO2/3 signal by immunofluorescence for GFP-positive and -negative cells from three independent experiments (Figure 1B). While GFP-PIAS3, -PIAS1, -ZNF451 and -LAZSUL reduced the average SUMO2/3 nuclear signal by 21%, 53%, 80% and 97%, respectively, GFP-PIAS4 positive cells presented a slight increase of 7% in SUMO2/3 signal. In a previous screen for targets of the SUMO Targeted Ubiquitin Ligase (STUbL) RNF4, we had observed that SUMO E3s were targets of RNF4 for ubiquitination and subsequent degradation by the proteasome, with ZNF451 and PIAS1 being the strongest RNF4 ubiquitination targets, and PIAS4 the weakest ^14, 15^. We hypothesized that overexpression of these E3s was promoting their hyperactivation leading to their auto-sumoylation and increased sumoylation of their substrates and subsequent degradation in an RNF4-dependent manner. To test our hypothesis, we made stable inducible U2OS cells for GFP-LAZSUL, which was the E3 with the strongest phenotype (Figure 1A-B), treated the cells with a control or an RNF4-targetting siRNA, induced the expression of the GFP-LAZSUL construct and analyzed the cells by immunostaining (Figure 1C-D) and immunoblotting (Figure 1E). RNF4 knock down caused an increase in the fraction of GFP-LAZSUL positive cells and rescued the SUMO2/3 depletion phenotype. Consistently, RNF4 knockdown increased the levels of both modified and non-modified GFP-LAZSUL.

**Figure 1.**
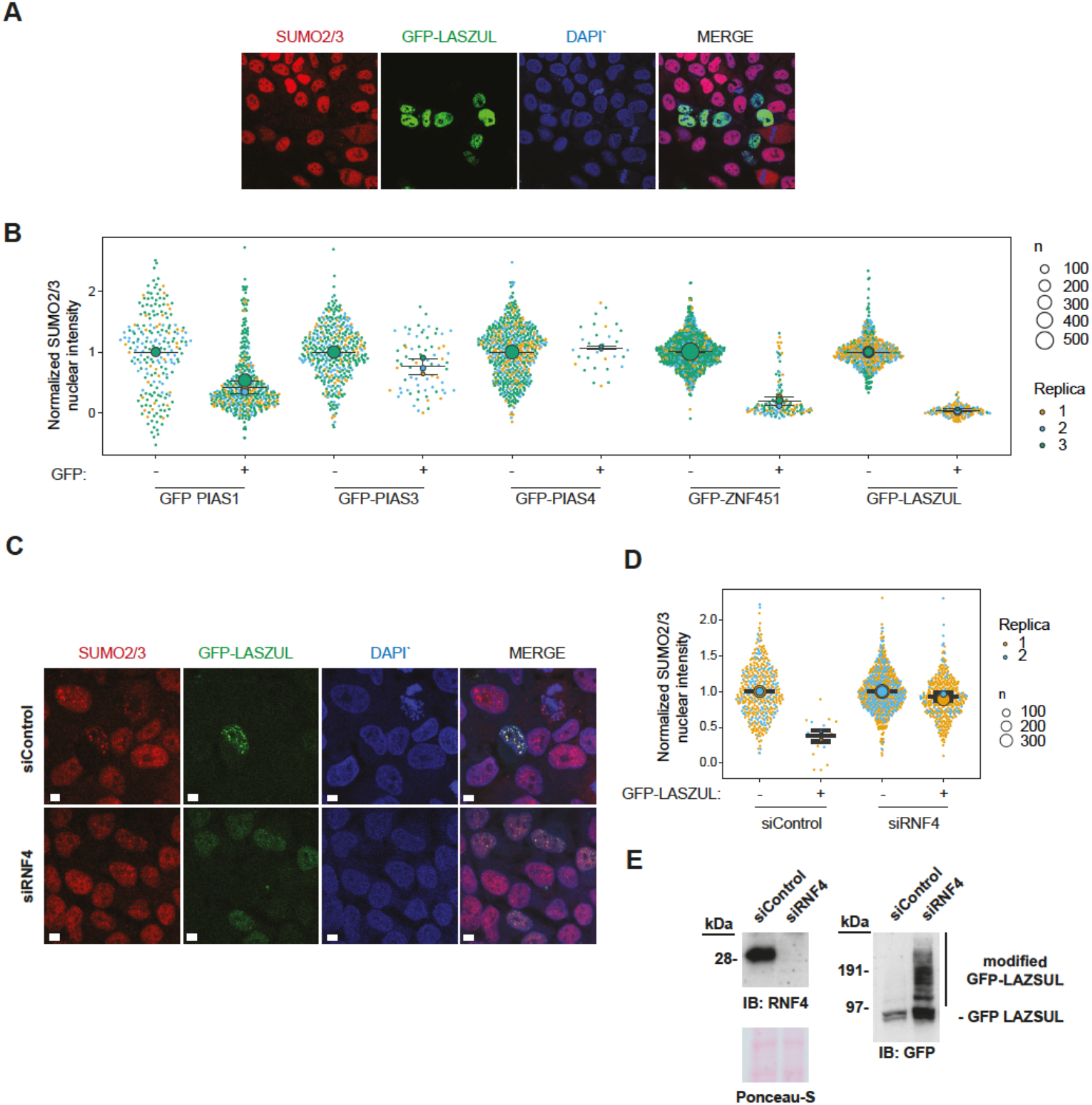
(A) Superplot depicting relative SUMO2/3 nuclear intensities after immunostaining of individual U2OS cells transiently transfected with GFP-tagged constructs of different E3s. Values were normalized to the average SUMO2/3 nuclear intensity of GFP-negatives from each individual experiment. Values from 3 independent experiments are depicted. (B) Representative immunofluorescence image of U2OS cells transiently transfected with GFP-LAZSUL immunostained for SUMO2/3. (C) Stable inducible GFP-LAZSUL expressing U2OS cells were treated with control or RNF4-targeting siRNAs and analyzed by immunostaining. (D) Quantification of the normalized nuclear SUMO2/3 intensities from the cells in (C). Independent values from two independent experiments are depicted. (E) Analysis by immunoblotting of the cells in (C). Size bars in fluorescence microscopy images represent 10 µm.

We conclude that SUMO E3 overexpression-based screens to identify SUMOylation substrates for specific SUMO ligases must be carefully evaluated since their conclusions could potentially be misleading because a negative control loop mediated by RNF4 is activated upon SUMO E3 overexpression, which might lead to SUMO2/3 depletion in cells.

### SUMO Activated Target Traps (SATTs) to identify the E3-specific SUMO proteome

Previously, in an effort to identify E3-specific ubiquitin substrates, Ubiquitin Activated Interaction Traps (UbAITs) were engineered^12^, which we later adopted and optimized for systematic screening in the Targets for Ubiquitin Ligases Identified by Proteomics (TULIP) methodology ^14, 15^. However, due to the high number of Ubiquitin E3 enzymes in the human proteome, performing the TULIP methodology on each E3 is an incredibly challenging task.

In contrast to Ubiquitin, the number of *bona fide* SUMO E3 enzymes is more limited, comprising the Siz/Pias Really Interesting New Gene (S-P RING) family, the ZNF451 family and RANBP2 ^1^. Therefore, addressing the E3-substrate wiring for SUMO E3s is a more manageable challenge.

Thus, similar to the TULIP2 methodology ^14^, we designed the SUMO Activated Target Traps (SATTs) approach, in which lentiviral doxycycline-inducible plasmids consisting of 10xHIS tag and a Gateway cloning sequence, followed by 10xHIS and either mature SUMO1 or mature SUMO2-Q87R were constructed. (Figure 2A) The Gateway sequence enables the straightforward shuttling of any SUMO E3 of interest. The SUMO2-Q87R mutation facilitates the identification of SUMO acceptor sites by mass spectrometry-based proteomics ^6, 7^. Consistently, the rationale behind this approach is that if we generate a linear fusion between an E3 and activated SUMO, the E3 will use the attached SUMO moiety to modify its substrate, enabling the co-purification of the E3 together with its substrate and subsequent identification by mass spectrometry-based proteomics. In line with TULIP2 methodology ^14^, we included two different negative controls in our screens. The first control is a ΔGG construct where the SUMO moiety lacks the C-terminal di-Gly motif and thus cannot be conjugated to a substrate. The second control is a catalytic dead mutant where the interaction with the SUMO E2 enzyme is abolished, thus the transfer of the SUMO moiety from the E2 to the substrate cannot be catalyzed (Figure 2B).

**Figure 2.**
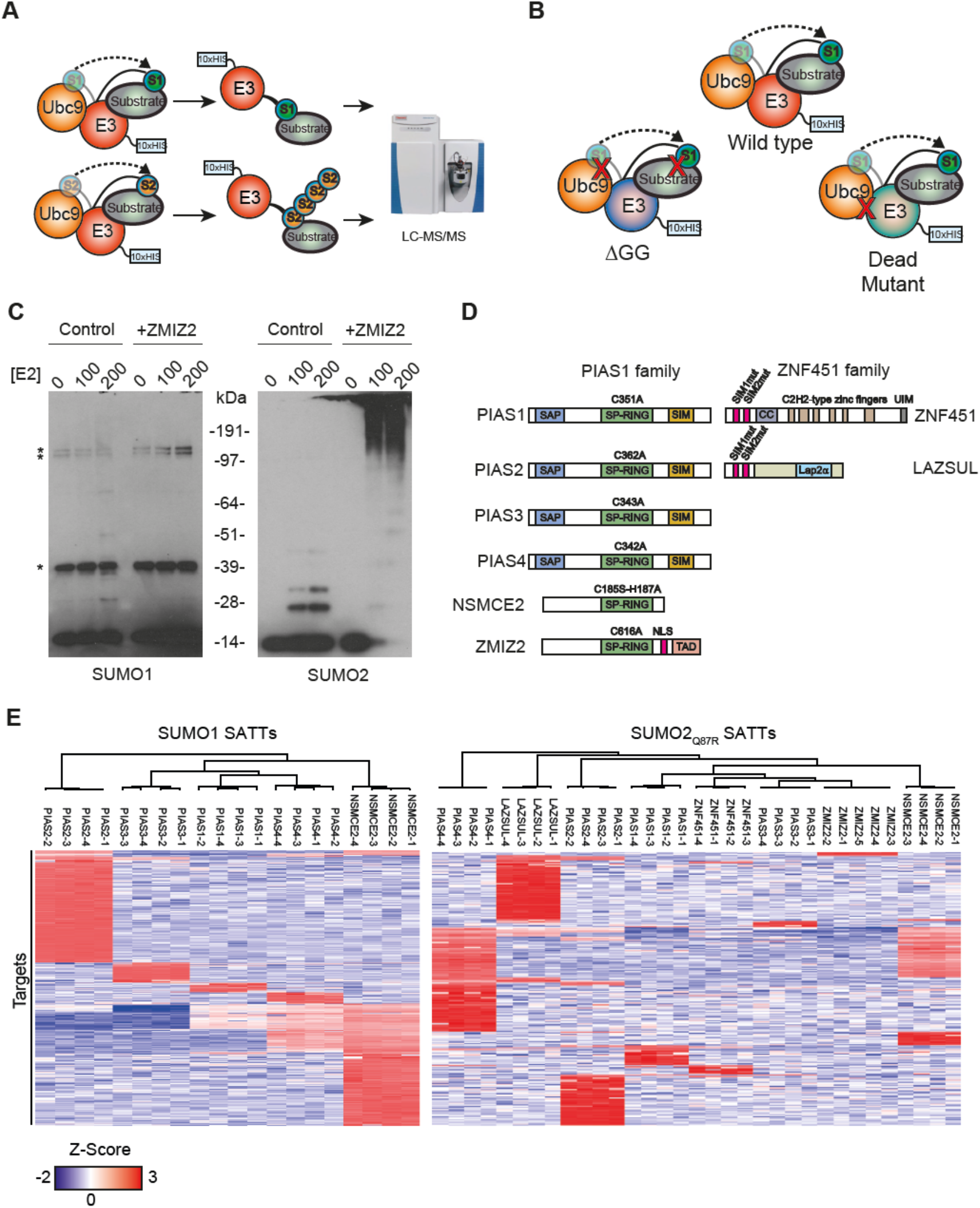
(A) SATTs screen rationale. SUMO moieties covalently attach to the C term of an E3 of interest will be attached to E3 substrates, enabling the co-purification of the E3 together with the sumoylation target, which will be later identified by mass spectrometry. (B) SATTs negative controls rationale. While ΔGG SATTs lack the C-terminal SUMO diGly motif, not enabling conjugation to the substrate, catalytic-dead mutants prevent interaction with the SUMO E2. (C) In vitro sumoylation assays including or ZMIZ2 SUMO E3 enzyme and different concentrations of the SUMO E2. Assays were carried out using either SUMO1 or SUMO2. (D) E3s studied in this article. The mutations performed on each E3 to construct the catalytic-dead mutant controls are indicated. (E) Heatmap depicting sumoylation targets for each of the replicates of the different E3s.

Furthermore, we tested *in vitro* the SUMO E3 activity of another PIAS-like enzyme, ZMIZ2^16–19^, for SUMO1 and SUMO2 ligase activity (Figure 2C) and observed E3 enzymatic activity for SUMO2 but not for SUMO1. Accordingly, we built SUMO1 SATTs for the S-P RING SUMO E3 enzymes, PIAS1, PIAS2, PIAS3, PIAS4 and NSMCE2. Additionally, for SUMO2 SATTs, we also included ZNF451, LAZSUL and ZMIZ2 as they are exclusive for SUMO2/3^20, 21^ (Supplementary Figure 2A). RANBP2 was left out from our screen due to the size of the protein (3,224 amino acids). Also, the ZNF451 family E3 KIAA1586 was left out because it is exclusively found in primates ^21^. To generate the catalytic-dead mutant controls, we introduced specific mutations in each E3 (Figure 2D). For the S-P RING family E3s, we mutated cysteines in the S-P RING domain, for the ZNF451 family E3s, we mutated the SUMO Interaction Motifs (SIMs) by substituting the long hydrophobic amino acids into alanines.

Next, we constructed U2OS cells stably expressing the inducible E3 SATTs constructs indicated in Supplementary Figure 2A, including the ΔGG and catalytic-dead mutant negative controls. We induced the expression of the constructs for 24h, lysed the cells in denaturing conditions and purified the SATT conjugates from four independent biological repeats (five in the case of ZMIZ2) yielding a total of 159 samples. In order to avoid RNF4-mediated degradation of the SATTs due to auto-sumoylation (Figure 1), and to increase the number of sumoylation conjugates ^7^, the proteasome was inhibited for 5h with MG132.

Sample analysis by immunoblotting (Supplementary Figure 2B-C) showed that the expression levels of the SATTs were below or close to endogenous counterparts for every construct Moreover, signal could be observed in a higher molecular weight smear for the wild type and catalytic-dead mutant construct for every SATT corresponding to E3-SUMO-target conjugates. This smear was absent in the ΔGG constructs. Consistently, the catalytic-dead SATT smears had different profiles than their wild type counterparts, indicating that the SUMO moieties in the mutant SATTs could still be used for conjugation by other endogenous E3s.

Mass spectrometry analysis of the samples identified 590 SUMO1 SATT conjugates and 1195 SUMO2 SATT conjugates, which were preferential or specific for the different E3s (Figure 2E, Supplementary Datasets 1-2).

### The SATT Index

Although the substrates we identified for each tested SUMO E3 were relatively specific for every E3 when comparing a wild type SATT with its ΔGG counterpart, all the substrates did not remain equally significant when comparing with their catalytic dead mutant counterpart (Supplementary dataset 1-2), indicating that, as previously shown for UbAITs ^12, 14, 15^, the SUMO moiety attached to the mutant SATT can also be conjugated to a substrate by another endogenous SUMO E3.

Therefore, we used the relation between the differences of the enrichment of a substrate for a specific E3 comparing both the ΔGG and the mutant counterpart to wildtype, which we termed SATT index:

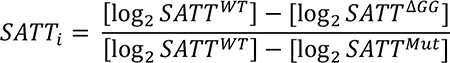

Values close to 1 and higher are considered very specific and values close to 0 and lower are considered not specific.

### Different E3s have different preferences towards SUMO1 or SUMO2/3

It could be argued that making a SUMO1 SATT with an E3 which normally catalyzes SUMO2/3 conjugation might force SUMO1 conjugation on SUMO2/3 substrate. Thus, we decided to investigate if SUMO E3s could discriminate substrate specificity depending on the SUMO type they were conjugating. For this purpose, we looked at the overlap between SUMO1 and SUMO2 substrates for the different S-P RING E3s that had been investigated both for SUMO1 and SUMO2 SATTs (Figure 3A). On one side, PIAS1 and PIAS3 had a very specific distinction between SUMO1 and SUMO2/3 substrates. For PIAS1, only 12% of the SUMO1 substrates were also SUMO2/3 substrates, and 9% of the SUMO2/3 were also substrates for SUMO1. For PIAS3 the numbers were 4% and 3% respectively. On the other side, PIAS4 had a very high preference towards SUMO2/3 conjugation and 88% of the SUMO1 substrates were also SUMO2 substrates, being the number of SUMO2/3 substrates 3.5 times higher, which indicates that PIAS4 is mainly a SUMO2/3 E3 enzyme as previously described^11^. In between PIAS2 and NSMCE2 had intermediate overlap percentages, which indicates an intermediate specificity for SUMO1 or SUMO2/3 substrates.

**Figure 3.**
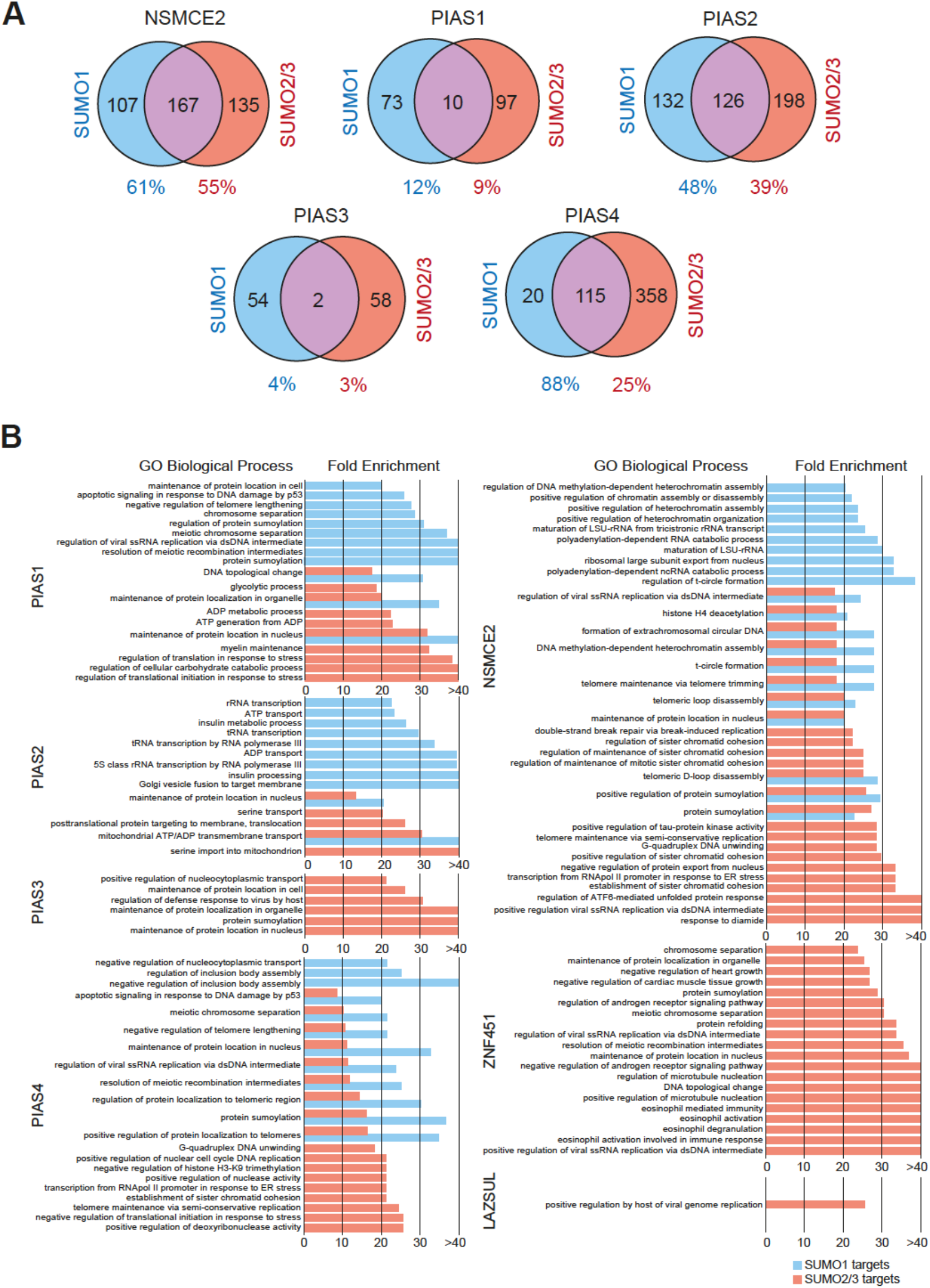
(A) Overlap between SUMO1 and SUMO2/3 SATT substrates for the indicated E3s. (B) Gene ontology analysis for the sumoylation substrates of the different E3 SATTs analysed in this study.

### Gene Ontology analysis

Gene Ontology analysis for biological processes of the sumoylation substrates for the different E3s indicated that different E3s are involved in different processes (Figure 3B, Supplementary Dataset 3). As expected, PIAS1, PIAS4, NSCME2 and ZNF451 substrates are enriched in Gene Ontology terms relative to genome biology ^22–28^, and PIAS3 substrates are enriched for maintenance of proteins at the nucleus ^29–31^. LAZSUL substrates were only mildly enriched for host viral genome replication term and no specific enrichment was present for ZMIZ2. Interestingly, PIAS2 substrates are enriched for membrane translocation and ADP/ATP mitochondrial transport and ZNF451 for immune-related processes, two processes to which these E3 enzymes have not been previously connected.

### PIAS4 and NSMCE2 make hybrid SUMO1-SUMO2/3 chains

SUMO2_Q87R_ SATTs leave a QQTGG remnant after tryptic digestion on acceptor lysines which can be identified by mass spectrometry-based proteomics ^7^. Although K11 is known to be the canonical site to make SUMO2/3 chains ^32^, several other SUMO2/3 sites at the endogenous level have been identified ^5^. Therefore, in addition to K11-SUMO2/3 chains, other chain types exist.

Mass spectrometry analysis of our samples enabled us to obtain MS/MS spectra in which the QQTGG remnant could be localized on SUMOs in an unambiguous manner (Figure 4A) and the intensity of these sumoylation sites could be quantified (Figure 4B). SUMO2/3 K11 chains, were found with every E3. As expected, no QQTGG-modified peptides were found in ΔGG SATTs samples. In contrast, signal for the modification with SUMO2/3 on K11 either on SUMO2 or SUMO3 could be detected in every SATT and, at less intense level, on K5. This included both wild type and catalytic-dead mutant SATTs.

**Figure 4.**
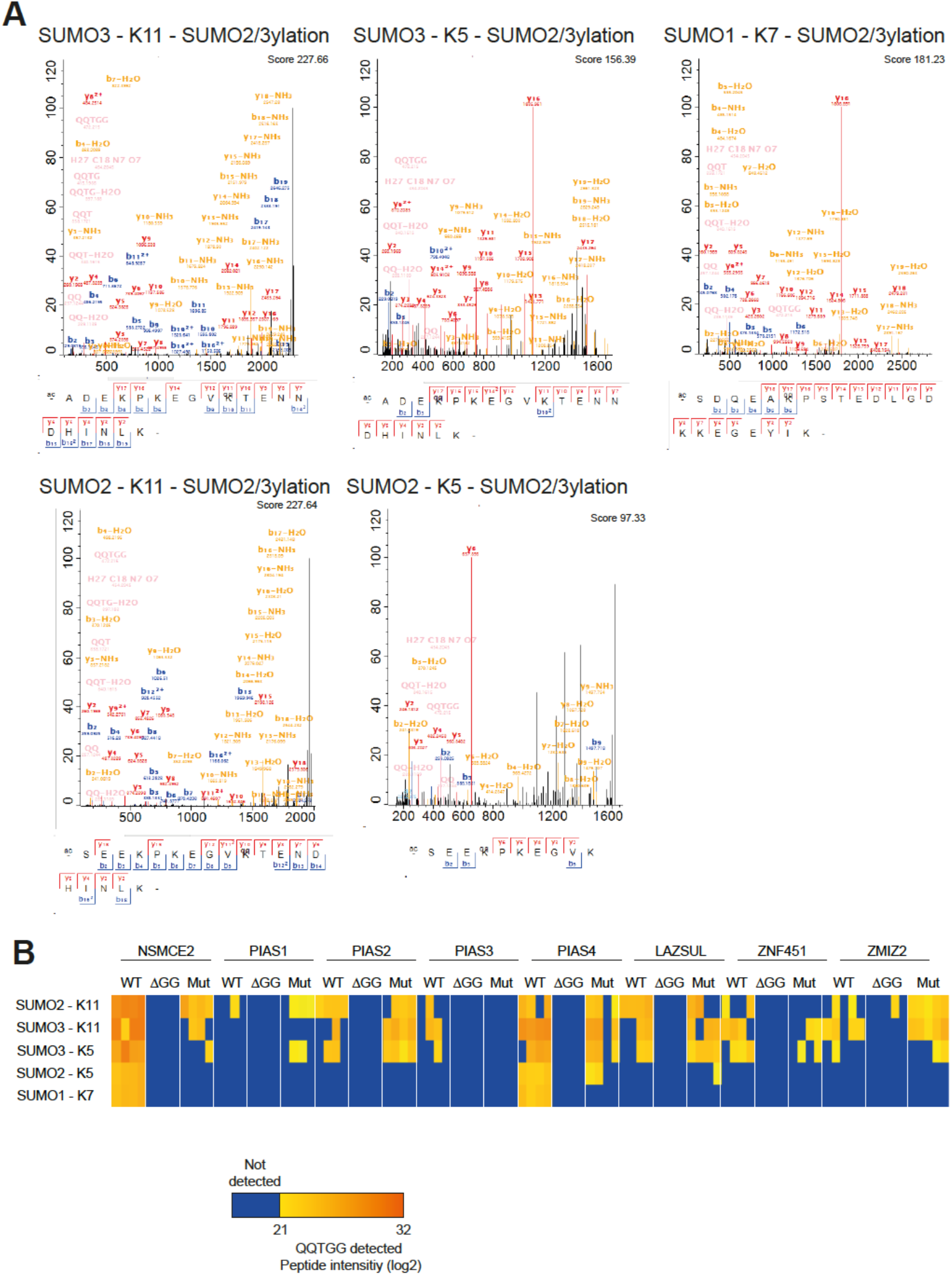
(A) Best MS/MS spectra corresponding to the QQTGG sites on SUMOs identified in an unambiguous manner. (B) Site intensities of the different sites depicted in (A) in the different SUMO2Q87R SATT samples.

Importantly, only NSCME2 and PIAS4 wild type SATTs were able to modify SUMO1 with SUMO2/3 at K7, being completely dependent on the catalytic activity of the SATT.

### Target validation - histone sumoylation

Among the 1681 E3-specific substrates that we had identified for both SUMO1 and SUMO2/3, we decided to focus on the E3 enzymes that were modifying histones with SUMO2/3 (Figure 5A). We found that LAZSUL was responsible for modifying histone H1, specifically H1.0, H1.2 and H1.4/H1.5 (identified peptides did not allow distinguishing between H1.4 and H1.5) types.

**Figure 5.**
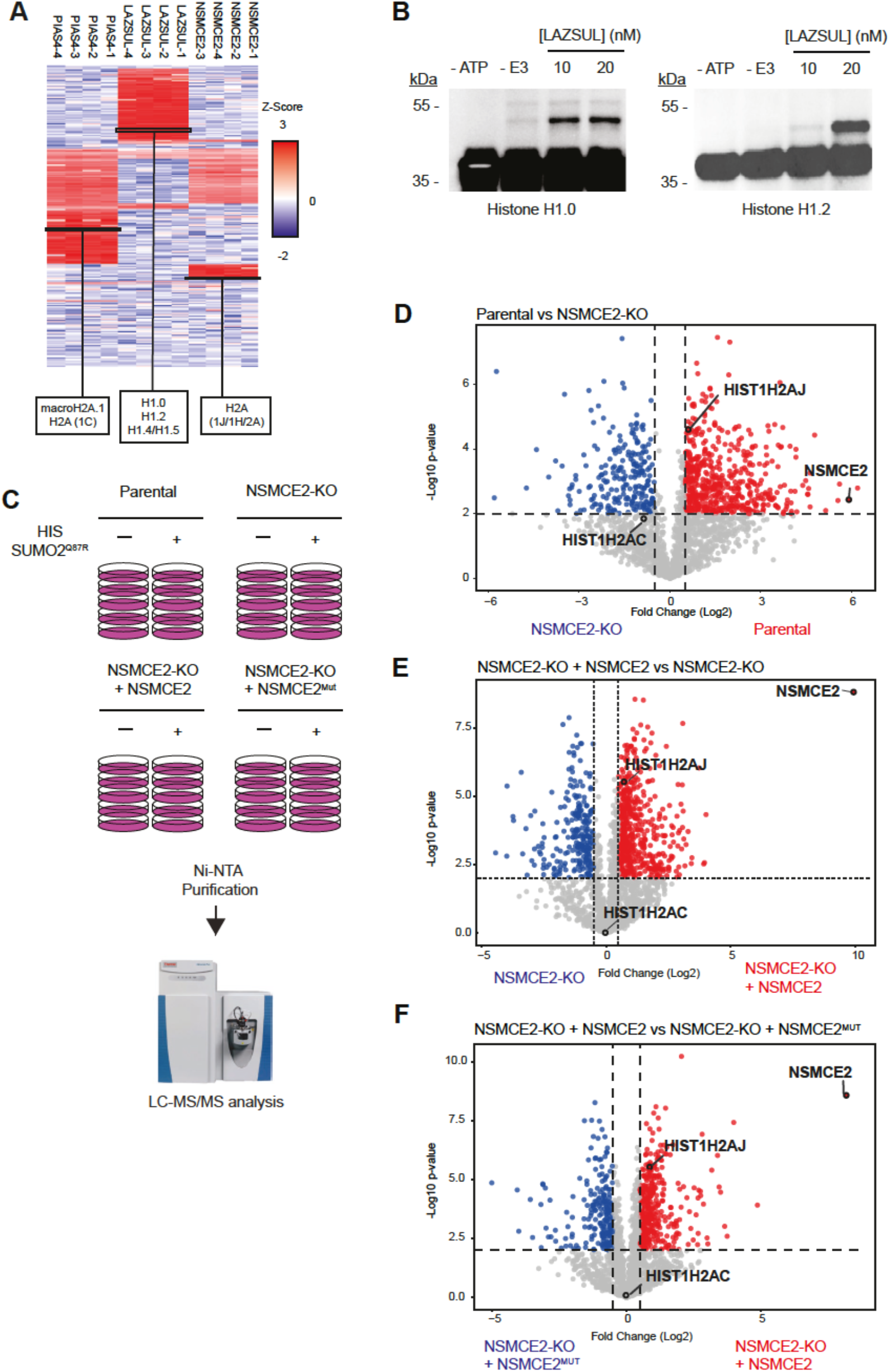
(A) Heatmap depicting relative enrichment of each of the identified sumoylation targets identified in each replicate for SUMO2Q87R SATTs which histones are identified as substrates. Corresponding histones are indicated. (B) . Inmunoblotting against histone H1.0 and H1.2 of in vitro sumoylation assays adding different concentrations of LAZSUL as E3. (C) Experimental setup. Different U2OS cell lines expression HIS-SUMO2Q87R where cultured in large scale, HIS-SUMO2Q87R conjugates purified using Ni-NTA beads and samples subsequently analyzed by mass spectrometry.

To test if LAZSUL was able to promote histone H1 sumoylation, we performed *in vitro* sumoylation assays with recombinant histone H1.0 and H1.2 (Figure 5B). In line with our results, LAZSUL was able to promote histone H1 sumoylation in a concentration-dependent manner.

NSMCE2 and PIAS4 shared a high number of sumoylation substrates, which mainly belonged to DNA Damage Response-related pathways (Figure 2E, 3B, Suppementary Dataset 1,2,3). Interestingly, PIAS4 and NSMCE2 had specificity for different histone H2A isoforms. While PIAS4 had histone macroH2A.1 and H2A type 1C as substrates, NSCME2 had histone H2A type 1J or other histone H2A types that we could not discriminate based on the identified peptides (Figure 5A). Our results indicate that the E3-to-substrate wiring was highly specific (Figure 2E, Suppementary Dataset 1,2). In the case of histone H2A this was very striking, where E3 specificity reached even the isoform level (Figure 5A). This specificity contrasts with the current paradigm in the field that SUMO E3s are highly redundant ^33^.

In contrast to LASZUL, which we used for *in vitro* sumoylation assays, NSCME2 belongs to the SMC5/6 complex, and its sumoylation activity is DNA-dependent^23^. Moreover, Histone H2A is incorporated into nucleosomes together with Histones H3, H4 and H2B. Other PTMs occur on other histones and they can co-exist and be mutually dependent. Thus, performing *in vitro* sumoylation assays with H2A and different E3s in relevant settings is much more challenging. Therefore, we aimed to test histone H2A to SUMO E3 specificity with an alternative approach. According to our results, deficiencies in NSMCE2 should affect histone H2A sumoylation of all types except for histone H2A type 1C.

We stably expressed or not HIS-tagged SUMO2 in parental and NSMCE2 knockout cells and rescued NSMCE2 knockout cells either with wild type NSMCE2 or with a catalytic dead mutant NSMCE2^C185S/H187A^^34^. We included a Q87R mutation in our HIS-SUMO2 construct to enable the identification of the sumoylation site on modified proteins by mass spectrometry-based proteomics^7^. Next, we cultured our different cell lines, lysed them, purified the HIS-SUMO2_Q87R_ proteome and identified substrates by mass spectrometry (Figure 5C). Quantitative proteomic analysis confirmed that the sumoylation of histone H2A-1J (HIST1H2AJ) required the catalytic activity of NSMCE2, whereas the sumoylation of histone H2A-1C (HIST1H2AC) was not affected (Figure 5D-F). These results confirm the results from our SATT screen.

### Polar SATTs – A user-friendly site to browse the dataset

Most proteomic screens, including this one, usually consist of large spreadsheet datasets full of gene/protein names and values and comparisons. The interpretation of these datasets can be daunting for researchers from other disciplines. To overcome these potential outreach hurdles, we developed an online web app tool to browse the dataset, which is freely accessible (https://amsterdamstudygroup.shinyapps.io/PolaRVolcaNoseR/). This tool enables users to select a protein of interest, and, if present in this study, will pop up in a polar plot in the sectors corresponding to the relevant E3s, indicating enrichment in terms of p-value and difference between wild type and ΔGG SATTs, and how relatively specific the substrate is for the E3 in terms of SATTi.

Moreover, the app can be used to customize the data visualization by enabling adjustment of the p-values, differences and SATTi limits, choosing to hide the values that exceed the limits. The size of the datapoints can also be adjusted to facilitate visualization, and the resulting figure highlighting the desired substrates can be exported in .pdf or .png. A “Dark Theme” is also available.

Finally, If users prefer browsing in independent Volcano Plots instead of the default Polar Plot, that is also enabled. An example of visualization of different sumoylation substrates for different E3s using the app is shown in (Figure 6)

**Figure 6.**
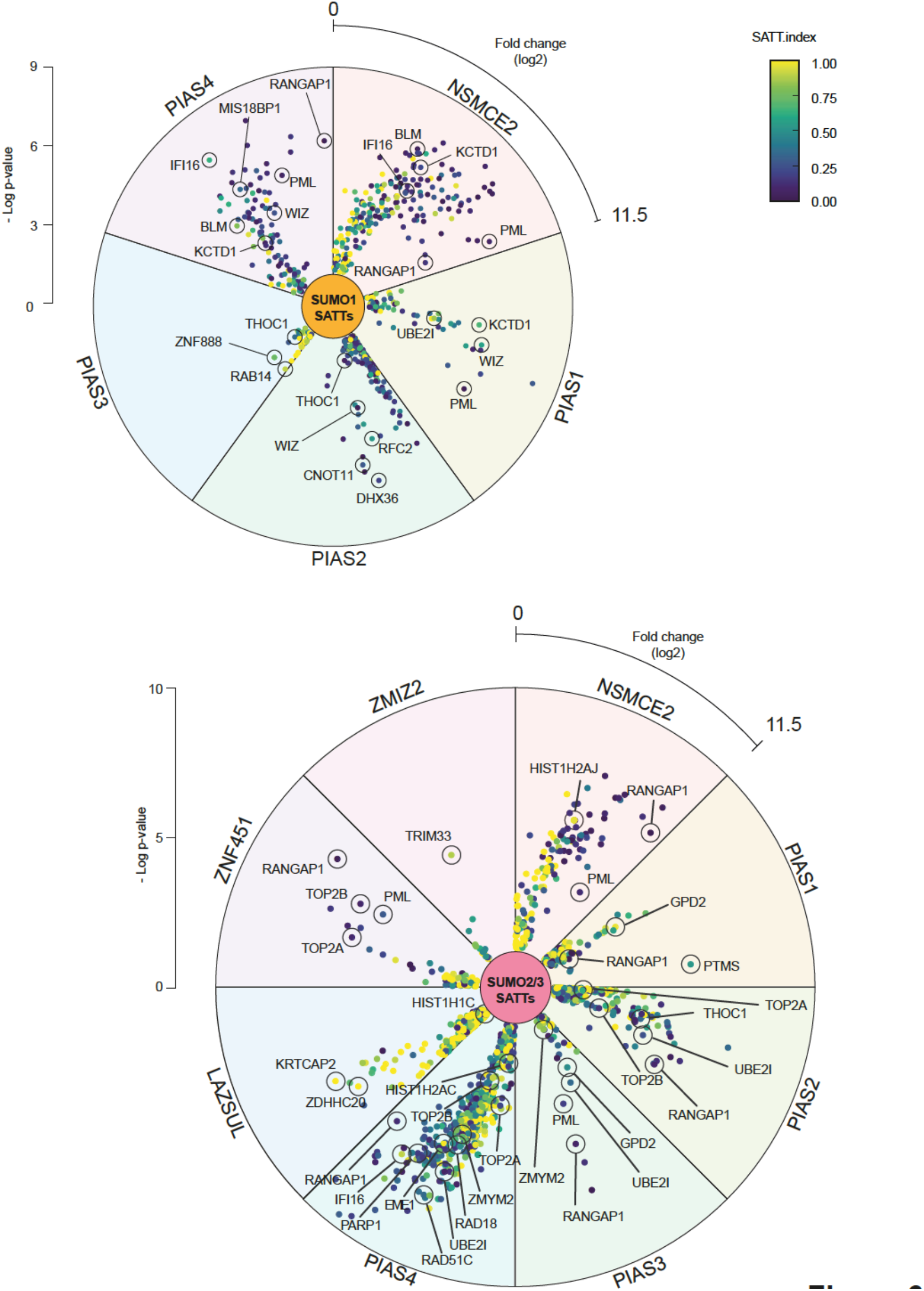
SATT Polar plots extracted from the polarVolcaNoseR web app. Some of the most prominent or specific substrates from different E3s are indicated.

## DISCUSSION

This resource identifies 590 SUMO1 and 1195 SUMO2/3 targets in an E3-specific manner. However, the number of sumoylation substrates that have been identified for SUMO1 and SUMO2 in U2OS cells is much higher ^6, 8, 9^. It is noteworthy, that the screens performed in this project only comprised a single condition of 5h of proteasome inhibition with MG132 in a single cell line, U2OS, for 8 different SUMO ligases.

The results obtained from this screen also corroborate previous observations on protein arrays regarding E3 preferences for SUMO1 or SUMO2/3^11^. PIAS3 and PIAS4 had been proposed to have a big preference for SUMO2. Consistently, for PIAS3, enrichments are much higher in SUMO2 SATTs than in SUMO1 SATTs (Figure 6), and, for PIAS4, 88% of SUMO1 substrates are shared with SUMO2. PIAS1 was observed to have no preference, in line with our SATT screen. For PIAS2 and NSMCE2 there is no data available from protein arrays to make this comparison.

Additionally, substrates that are highly sumoylated shared by every E3, such as PML, RANGAP, RNF216 or ZBTB33 have a very low SATT index for every E3, which indicates that either they are strong substrates for other E3s not included in this screen or require no E3 at all for sumoylation. In contrast, SATTi for histones were much higher (Supplementary Datasets 1-2). Moreover, previously independently described substrates for specific E3s had the best SATTi value for the relevant E3. Namely, PARP1^35^, ZMYM2^36^, and TOP2a^24, 27^ for PIAS4 and SUMO2/3 or histone H2A except type 1C for NSMCE2 (Figure 5).

### SUMO2/3 is not recycled at the proteasome

Whereas the classical model suggests that SUMO2/3 is deconjugated and recycled at the proteasome ^37^, we have observed that when overexpressing PIAS1, ZNF451 or LAZSUL, SUMO2/3 is depleted from the nucleus in an RNF4-dependent manner (Figure 1). This indicates that SUMO2/3 moieties attached to STUbL targets are also degraded. Whether the proteasome discriminates between mixed SUMO2/3-Ubiquitin chains on substrates and substrates that are co-modified independently by SUMO2/3 and Ubiquitin chains requires further investigation. We previously showed that the oncogene c-Myc, which sumoylated form highly accumulates upon proteasome inhibition and is and RNF4 target ^7, 14, 38^, is sumoylated and ubiquitinated on different residues in an RNF4-dependent manner. Based on these results, we favour the hypothesis that independent ubiquitination of the sumoylated substrate is sufficient for the degradation of SUMO moieties attached to the substrate without the need for mixed chains. Nevertheless, we also found that mixed SUMO/ubiquitin polymers are efficiently stabilized by proteasome inhibitors, indicating that these mixed polymers also constitute an efficient proteasomal degradation signal ^4^.

In contrast to GFP-PIAS1, -ZNF451, -LAZSUL and, partially, -PIAS3, which overexpression caused nuclear SUMO2/3 depletion, GFP-PIAS4 overexpression was linked to a slight increase in nuclear SUMO2/3 levels (Figure 1). Interestingly, PIAS4 and NSMCE2 were the only E3s capable of assembling SUMO2/3-SUMO1 mixed chains (Figure 4). We therefore hypothesize that SUMO2/3-SUMO1 mixed chains are poor substrates for RNF4. Similarly, SUMO1-capped SUMO2/3 chains are poor substrates for RNF4 but efficient substrates for RNF111/Arkadia, another STUbL^39^. Previously, it was shown that both SUMO1 and SUMO2 were recruited to sites of DNA damage, with SUMO1 recruitment depending on PIAS4^22^. Accordingly, DNA damage repair pathways are significantly enriched among PIAS4 substrates for SUMO1 (Figure 3B). However, we did not detect affinity of DNA Damage Response-related proteins for SUMO1 moieties ^9^. Which, might indicate that SUMO1 moieties incorporated at DNA damage sites in a PIAS4-dependent manner correspond to hybrid SUMO1-SUMO2/3 chains. Our results indicate that different types of mixed SUMO polymers and mixed SUMO/ubiquitin polymers constitute differential signals.

### Perspective and implications

Sumoylation of proteins occur in response to many different types of cellular stresses, such as DNA damage or replication stress and heat shock among others ^28, 40^. The E3 that modifies a specific target may vary depending on the scenario, and SATTs should be screened for specific targets at specific conditions. One E3 can be specific for the sumoylation of a protein in a certain context, and another in a different context. In this regard, PIAS4 and NSCME2 shared many sumoylation substrates from the DNA damage response. We speculate that, while NSMCE2 sumoylates these substrates in response to DNA damage in the context of DNA replication as part of the SMC5/6 complex^41^, PIAS4 sumoylates these substrates in a replication-independent manner.

Nonetheless, results from this screen might open new lines of investigation. Gene ontology analysis revealed that PIAS2 substrates are significantly enriched for serine import into mitochondrion (Figure 3B) as biological function, which deregulation has been very recently described as causative for Parkinson disease ^42^. Interestingly, PIAS2 has also been recently linked to Parkinsonism^43^. This suggest a potential role for the identified PIAS2 substrates in the development of this neurological disorder. Furthermore, ZNF451 substrates are very significantly enriched for eosinophil activation biological functions, which also correlates with the high expression of ZNF451 in bone marrow ^44^. Future mining of the resource presented here could improve our understanding of the biological functions of the different SUMO E3 ligases.

## METHODS

### Antibodies

Antibodies are listed in Table 1 with working dilutions.

### Generation of SATT toolbox

For the generation of the SATTs plasmids AgeI-10HIS-SUMO1-XmaI, AgeI-10HIS-SUMO1ΔGG-XmaI, AgeI-10HIS-SUMO2_Q87R_-XmaI, AgeI-10HIS-SUMO1ΔGG_Q87R_-XmaI restriction fragments from PCR products amplified with primers FW-AgeI-10HIS-SUMO1, FW-AgeI-10HIS-SUMO2, RV-XmaI-SUMO1, RV-XmaI-SUMO1noGlyGly, RV-XmaI-SUMO2-Q87R and RV-XmaI-SUMO2-Q87R-noGlyGly, were cloned into AgeI-SpeI sites of pCW57.1-nonStop plasmid^15^. The N-terminal 10xHIS tag was cloned as previously done for the H-TULIP2 plasmids^14^. Primer sequences are listed in Table T2.

### Generation of SATT lentiviral plasmids

SUMO1 and SUMO2 SATT plasmids containing a SUMO E3 ligase of interest were generated using Gateway® cloning LR reaction (Thermo Fisher Scientific). LR reactions were performed using a donor plasmid containing an E3 enzyme cDNA without stop codon and a SATT plasmid as destination vector. We used different donor plasmids. pENTR223-PIAS1, pDNOR221-PIAS2, pENTR223-PIAS3, pDNOR221-PIAS4, pENTR223-NSMCE2, and pENTR233-ZMIZ2 were obtained from DNASU repository with the following IDs: HsCD00505402, HsCD00042133, HsCD00514170, HsCD00041383, HsCD00287670 and HsCD00505806 respectively. pDNOR207-ZNF451-1 (isoform 1) and pDNOR207-ZNF451-3 (isoform 3) were generated using the Gateway® cloning BP reaction (Thermo Fisher Scientific) upon cDNA amplification using BP-tailed primers and pDNOR207 as donor vector. Catalytic dead mutants of each SUMO E3 ligase were generated by site-directed mutagenesis. Primer sequences are listed in Table T2.

### Transfection

For the transient transfection experiments in Figure 1, 10^4^ U2OS cells were transferred to 6-well plates containing 18 mm coverslips and left to attach overnight. The next day, 300 µL of transfection mixture consisting of 150 mM NaCl containing 1 µg of plasmid DNA and 6µg of Polyethylenimine were added to the cells. 24h after transfection, culture medium was replaced for fresh medium. Cells were fixed 48h after transfection for immunofluorescence analysis.

### siRNA for RNF4 transfection

siRNA-mediated knockdowns were performed as previously described^15^. In brief, DharmaFect 1 Transfection Reagent (GE Lifesciences) was used, according to the manufacturer’s instructions using on-target plus RNF4 siRNAs J-006557-08 and the non-targeted control was performed using siGENOME Non-Targeting siRNA #1 (GE Lifesciences).

### Cell culture

293T HEK, U2OS and RPE1 cells were cultured in Dulbecco’s modified Eagle’s medium (DMEM) supplemented with 10 % Fetal Bovine Serum (FBS) and 100 U/mL penicillin/100 µg/mL streptomycin at 37°C and 5% CO2 unless specified. Cells were regularly tested for mycoplasma contamination.

### SATT lentivirus production

293T HEK cells were seeded at 30% confluency in a T175 flask containing 16 mL of DMEM + 10% FBS. The following day, a 2 mL transfection mixture containing lentiviral packaging plasmids 7.5 µg pMD2.G (#12259, Addgene), 11.4 µg pMDLg-RRE (#12251, Addgene), 5.4 µg pRSV-REV (#12253, Addgene) and 13.7 µg SATT plasmid with 114 µL of 1 mg/mL Polyethylenimine (PEI) was prepared in 150 mM NaCl. After vortexing, solutions were incubated 10 min at room temperature before adding to the HEK cells. The day after transfection, culture medium was changed by fresh DMEM/FBS/Pen/Strep. 3 days after transfection, lentiviral suspension was harvested by filtering through a 0.45 µm syringe filter (PN4184, Pall Corporation). Lentiviral particle concentration was determined using the HIV Type 1 p24 antigen ELISA Kit (ZeptoMetrix Corporation).

### Generation of SATT and GFP-LAZSUL cell lines

U2OS cells were seeded in a 15cm diameter plates at 10% confluency with DMEM + 10% FBS. The next day, cell culture medium was replaced with lentiviral SATT constructs containing medium and polybrene 8 µg/mL. Lentiviral containing medium was replaced by fresh DMEM/FBS/Pen/Strep culture medium after 24h. After 2 days in fresh medium, 3 µg/mL puromycin was added to the medium for selection of SATT positive clones.

### Anti-SUMO2/3 immunostaining

Cells were grown on 9 mm coverslips and fixed with 4% paraformaldehyde for 15 minutes at 37°C in PBS and permeabilized with 0.1% Triton X-100 in PBS for 15 minutes. Next, cells were washed twice with PBS and once with PBS with 0.1% Tween-20 (PBS-T). Cells were then blocked for 10 minutes with 0.5% blocking reagent (Roche) in 0.1 M Tris, pH 7.5 and 0.15 M NaCl (TNB), and treated with anti-SUMO2/3 antibody in TNB for one hour. Coverslips were washed five times with PBS-T and incubated with the secondary antibody (Goat anti-mouse coupled to Alexa-594) in TNB for one hour. Next, coverslips were washed five times with PBS-T and dehydrated by washing once with 70% ethanol, once with 90% ethanol, and once with 100% ethanol. After drying the cells, coverslips were mounted onto a microscopy slide using citifluor/DAPI solution (500 ng/mL).

Immunofluorescence image analysis was performed using the FiJi – ImageJ distribution^45^.

### Generation of His-SUMO2-Q87R U2OS cell lines

U2OS NSMCE2-KO cell lines were kindly provided by Dr. Geert Hamer ^46^. Lentiviral plasmids pLX303 (Addgene #25897) containing either NSMCE2-WT or NSMCE2-C185S-H187A catalytic dead mutant were generated by Gateway® cloning LR reaction and used to generate stable U2OS cell lines upon 5 µg/mL blasticidin.

U2OS, U2OS NSMCE2-KO and U2OS NSE2-KO rescued with either NSMCE2-WT or NSMCE2-C185S-H187A catalytic dead cells were infected using a bicistronic lentivirus encoding 10xHis-SUMO2-Q87R-IRES-GFP separated by an IRES, which was modified from previously described 10xHis-SUMO2-WT ^47^. Following infection, U2OS cells were sorted in an FACSAria III (BD Biosciences) for low GFP levels.

### Purification of SATT conjugates

Following the TULIP2 methodology ^14^, five 15 cm diameter plates of U2OS cells expressing a particular SUMO E3 ligase, were grown up to 60% to 80% confluence. Expression of SATTs constructs was induced with 1µg/mL doxycycline once 60-80% confluence was reached. 24h after doxycycline induction, cells were treated with proteasome inhibitor MG132 (Sigma Aldrich) at 10 µM for 5h. After proteasome inhibition, cells were washed twice with ice-cold PBS and scraped. Cells were spun down and collected in 2mL ice-cold PBS, 100 µL of sample was taken as input and lysed in 200 µL SNTBS buffer (2% SDS, 1% NP-40, 50mM TRIS pH 7.5, 150 mM NaCl). After additional centrifugation, cells were lysed in 10mL Guanidinium buffer (6M guanidine-HCl, 0.1M Sodium Phosphate, 10mM TRIS, pH 7.8) and snap frozen in liquid nitrogen.

After thawing, lysates were homogenized at room temperature by sonication at 80% amplitude during 5s using a tip sonicator (Q125 Sonicator, QSonica, Newtown, USA). Sonication was performed twice. Subsequently, protein concentration was determined by BiCinchoninic Acid (BCA) Protein Assay Reagent (Thermo Scientific). Total protein in lysates was equalized accordingly. After equalization, lysates were supplemented with 5 mM β-mercaptoethanol and 50 mM Imidazole pH 8.0. 100 µL of dry nickel-nitrilotriacetic acid-agarose (Ni-NTA) beads (QIAGEN), were equilibrated with Guanidinium buffer supplemented with 5 mM β-mercaptoethanol and 50 mM Imidazole pH 8.0. Equilibrated Ni-NTA beads were added to the cell lysates and incubated overnight at 4°C under rotation.

After lysate-bead incubation, Ni-NTA beads were spun down and transferred with Wash Buffer 1 (6 M Guanidine-HCl, 0.1M Sodium Phosphate, 10 mM Tris, 10 mM Imidazole, 5 mM β-mercaptoethanol, 0.2 % Triton X-100, pH 7.8) to an Eppendorf LoBind tube (Eppendorf). After mixing and spinning down again, the beads were moved to a new LoBind tube with Wash buffer 2 (8 M Urea, 0.1M Sodium Phosphate, 10 mM Tris, 10 mM imidazole, 5mM β-mercaptoethanol, pH 8). This procedure was repeated with Wash buffer 3 (8 M urea, 0.1M Sodium Phosphate, 10 mM TRIS, 10 mM imidazole, 5 mM β-mercaptoethanol, pH 6.3). Ultimately, beads were washed twice with Wash buffer 4 (8 M urea, 0.1M Sodium Phosphate, 10 mM TRIS, 5 mM β-mercaptoethanol, pH 6.3). When washing with wash buffer 3 and 4, beads were allowed to equilibrate with the buffer for 15 min under rotation.

### Trypsin digestion of SATT-purified conjugates

After the final wash with Wash buffer 4, Ni-NTA beads were resuspended in 7 M urea, 0.1 M NaH2PO4/Na2HPO4, 0.01 M Tris/HCl, pH 7 and digested with 500 ng recombinant Lys-C (Promega) at RT while shaking at 1,400 rpm. After 5h with Lys-C, urea buffer was diluted to <2M by adding 50 mM ABC. A second digestion was performed o/n at 37°C while shaking at 1,400 rpm using 500 ng of sequencing grade modified trypsin (Promega). Trypsin digested peptides were separated from Ni-NTA beads by filtering through a 0.45 µm filter Ultrafree-MC-HV spin column (Merck-Millipore).

### Purification of 10xHis-SUMO2 conjugates

10xHis-SUMO2 and 10xHis-SUMO2-Q87R conjugates were purified using Ni-NTA beads as previously described ^48^. In brief, cells were grown in five 15 cm dishes and were or were not treated with 10µM proteasome inhibitor (MG132) for 5h. Subsequently, cells were scraped and lysed in 6M guanidinium buffer. A small fraction of cells was separately lysed in SNTBS buffer as input control. After homogenization by sonication, lysates were incubated with Ni-NTA beads o/n at 4 °C. The next day, Ni-NTA beads were washed with buffers 1-4 and eluted in 7 M urea, 0.1 M NaH2PO4/Na2HPO4, 0.01 M Tris/HCl, pH 7.0, 500 mM imidazole pH7.

### Trypsin digestion of SUMO2 purified conjugates

Eluted proteins were supplemented with ABC to 50 mM. Subsequently, samples were reduced with 1 mM dithiothreitol (DTT) for 30 min and alkylated with 5 mM chloroacetamide (CAA) for 30 min. After an additional reduction with 5 mM DTT for 30 min at RT, conjugates were digested with Lys-C for 5h at RT. After Lys-C digestion, peptides were diluted with 50 mM ABC and trypsin digested o/n in dark.

### Mass Spectrometry sample preparation

Digested peptides were acidified by adding 2% TriFlourAcetic (TFA) acid. Subsequently, peptides were desalted and concentrated on triple-disc C18 Stage-tips as previously described ^49^. Stage-tips were in-house assembled using 200 µL micro pipet tips and C18 matrix. Stage-tips were activated by passing through 100µL of methanol. Subsequently 100 µL of Buffer B (80% acetonitrile, 0.1% formic acid), 100 µL of Buffer A (0.1% formic acid), the acidified peptide sample, and two times 100 µL Buffer A were passed through the Stage-tip. Elution was performed in 50 µL of 32.5% acetonitrile, 0.1% formic acid.

Samples were vacuum dried using a SpeedVac RC10.10 (Jouan, France) and stored at −20°C. Prior to mass spectrometry analysis, samples were reconstituted in 10 µL 0.1% formic acid and transferred to autoload vials.

### LC-MS/MS analysis

All the experiments were analyzed by on-line C18 nanoHPLC MS/MS with a system consisting of an Ultimate3000nano gradient HPLC system (Thermo, Bremen, Germany), and an Exploris480 mass spectrometer (Thermo, Bremen, Germany). Samples were injected onto a cartridge precolumn (300 μm × 5 mm, C18 PepMap, 5 μm, 100 A) with a flow of 10 μl/min for 3 minutes (Thermo, Bremen, Germany) and eluted via a homemade analytical nano-HPLC column (50 cm × 75 μm; Reprosil-Pur C18-AQ 1.9 μm, 120 A) (Dr. Maisch, Ammerbuch, Germany). The gradient was run from 2% to 38% solvent B (80% acetonitrile, 0.1% formic acid) in 120 min. The nano-HPLC column was drawn to a tip of ∼10 μm and acted as the electrospray needle of the MS source. The temperature of the nano-HPLC column was set to 50°C (Sonation GmbH, Biberach, Germany). The mass spectrometer was operated in data-dependent MS/MS mode for a cycle time of 3 seconds, with a HCD collision energy at 28 V and recording of the MS2 spectrum in the orbitrap, with a quadrupole isolation width of 1.2 Da. In the master scan (MS1) the resolution was 120,000, the scan range 350-1600, at an standard AGC target with maximum fill time of 50 ms. A lock mass correction on the background ion m/z=445.12 was used. Precursors were dynamically excluded after n=1 with an exclusion duration of 45 s, and with a precursor range of 10 ppm. Charge states 2-5 were included. For MS2 the scan range mode was set to automated, and the MS2 scan resolution was 30,000 at an normalized AGC target of 100% with a maximum fill time of 60 ms.

### Mass Spectrometry data analysis

All raw data was analyzed using MaxQuant (version 1.6.14) as previously described ^50^ We performed the search against an in silico digested UniProt reference proteome for Homo sapiens including canonical and isoform sequences (5th July 2021). Database searches were performed according to standard settings with the following modifications: Digestion with Trypsin/P was used, allowing 4 missed cleavages. Oxidation (M), Acetyl (Protein N-term), Phospho (S,T) and QQTGG (K) (for SUMOylation sites) were allowed as variable modifications with a maximum number of 3. Carbamidomethyl (C) was disabled for SATTs analysis as a fixed modification. Label-Free Quantification was enabled, not allowing Fast LFQ. All peptides were used for protein quantification.

Output from the analysis in MaxQuant was further processed in the Perseus computational platform version 1.6.14 ^51^ for statistical analysis. LFQ values were log2 transformed and contaminants, proteins identified by site and reverse peptides were excluded by the analysis, as well as proteins that were not identified in the four replicates of at least one condition. Missing values were imputed from a normal distribution with a width of 0.3 and a downshift of 2.5. E3s were compared independently regarding wild type and ΔGG SATTs, and proteins that were not significantly enriched in the wild type SATTs were also excluded for the next comparison of the wild type versus the mutant SATTs. Finally, statistical analysis from every E3 SATT were combined again and missing values randomly inputed from a normal distribution with a width of 0.3 and a downshift of 2.5. Z-score was calculated for heatmap visualization. Resulting data were exported and further processed in Microsoft Excel 365 for comprehensive data visualization and calculation of SATT indexes.

Gene Ontology analyses were performed using the PANTHER overrepresentation test from the Gene Ontology Consortium^52^.

### Mass spectrometry data availability

The mass spectrometry proteomics data have been deposited to the ProteomeXchange Consortium via the PRIDE partner repository ^53^ with the dataset identifiers PXD034459.

For reviewing purposes, the following credentials can be used:

Username

Password: B3X8P566

### Electrophoresis and immunoblotting

Samples were separated on Novex 4-12% gradient gels (Thermo Fisher Scientific) using NuPAGE® MOPS SDS running buffer (50mM MOPS, 50mM Tris-base, 0.1% SDS, 1mM EDTA pH 7.7) and transferred onto Amersham Protran Premium 0.45 NC Nitrocellulose blotting membranes (GE Healthcare) using a Bolt Mini-Gel system (Thermo Fisher Scientific), which was used for both the gel electrophoresis and the protein transfer to the membrane according to vendor instructions. Membranes were stained with Ponceau-S (Sigma Aldrich) to determine total amount of protein loaded. Next, membranes were blocked with blocking solution (8% Elk milk, 0.1% Tween-20 in PBS) for 1h prior to primary antibody incubation. Chemiluminescence reaction was initiated with Western Bright Quantum Western blotting detection kit and measured in a ChemiDoc^TM^ imaging system (BIO-RAD, Hercules, CA, USA).

### In vitro SUMOylation

For the *in vitro* histone H1 sumoylation assays by LAZSUL, recombinant enzymes and substrate were used: E1: His-Aos1/His-Uba2; E2s, ZNF451 iso3; SUMO2 (Eisenhardt et al., 2015) and histone H1 (catalog # 81126) and H1.2 (catalog # 81252) from Active motif. All recombinant proteins are frozen in small single-use aliquots at the highest possible concentrations and stored at −80°C. Sumoylation assay buffer (SAB): 20mM HEPES pH 7.3, 110mM KOAc, 2mM Mg(OAc)2, 1mM DTT, 0.2mg/mL ovalbumin, 0.05%, Tween 20 (v/v). ATP: 100mM in 20mM HEPES pH 7.3, 100mM Mg(OAc)2, SDS sample buffer (3x): 150mM Tris–HCl (pH 6.8), 6% SDS, 30% glycerol, 0.3% bromophenol blue, 100mM DTT.

Briefly, an in vitro sumoylation reaction was set up in a final volume of 20μL by mixing together 60nM E1, 50 nM E2, 200 nM of the substrate (histone H1.0 and H1.2), and 2μM of SUMO2, and different concentrations of E3s (10, 20, 40, 80 nM and 160 nM for ZNF451 iso3). ATP was added to a final concentration of 5mM to start the reaction. Samples were incubated at 30°C for 30min and he reaction was stopped by adding 10μL 3x SDS sample buffer and heat denaturing for 5min at 95°C. 13μL of the sample was size-separated on a 15% SDS gel. Proteins were transferred onto nitrocellulose membranes by standard semidry protein transfer. The substrate and the SUMO2 modification were detected by immunoblotting using substrate-specific antibodies.

### Data Representation

Super plots and Volcano plots were constructed using SuperPlotsOfData^54^ and VolcanoseR ^55^ web apps.

For the SATT polar plots, a web app for the display of multiple volcano plots side-by-side, named polarVolcaNoseR, was made with R/Shiny.

The code was written using R (https://www.r-project.org) and Rstudio (https://www.rstudio.com). To run the app, several freely available packages are required: shiny, ggplot2, magrittr, dplyr, ggrepel, htmlwidgets, ggiraph, glue, scales. The web app is freely accessible at: https://amsterdamstudygroup.shinyapps.io/PolaRVolcaNoseR/ and the code is available at github: https://github.com/ScienceParkStudyGroup/polarVolcaNoseR. In the default ‘polar’ representation, the volcano plots are plotted in a circle, where the radius depicts the −Log_10_(p-value) and the circumference reflects the positive Log_2_(Fold-change). Labels of proteins can be added by point-and-click and the data of individual dots are displayed when the cursor hovers over a datapoint. A customized, interactive plot can be exported as a HTML file.

## AUTHOR CONTRIBUTIONS

RG-P and ACOV conceived the project. CvdM performed the GFP-tagged E3 overexpression experiments. DS-L prepared all the resource reagents and samples. AdR, AO and PvV injected mass spectrometry samples. DS-L and RG-P analyzed mass spectrometry data. EG made recombinant ZMIZ2. RG-P and EN performed *in vitro* sumoylation assays. JG programmed the polarVolcanoseR web App. AP supervised EN. RG-P supervised DS-L and CvdM. ACOV supervised EG. PvV supervised AdR and AO. RG-P lead the project and wrote the manuscript together with DS-L and input from all the authors.

## ACKNOWLEDGEMENTS

This work was supported by a Young Investigator Grant from the Dutch Cancer Society (KWF-KIG 11367/2017-2) and the EMERGIA program from the Andalusian Regional Government, Spain (Junta de Andalucia – EMERGIA20_00276) to RG-P. Work in the laboratory of A.C.O.V. has been supported by the European Research Council (ERC; grant 310913) and the Dutch Research Council (NWO; grant 724.016.003).

## FIGURE AND DATASET LEGENDS

**Supplementary Dataset 1.** SUMO1 SATTs dataset. Spreadsheet including identified SUMO1 targets for the different SATTs, SATTi, Statistic comparisons and miscellaneous MS/MS proteomics values.

**Supplementary Dataset 2.** SUMO2_Q87R_ SATTs dataset. Spreadsheet including identified SUMO1 targets for the different SATTs, SATTi, Statistic comparisons and miscellaneous MS/MS proteomics values.

**Supplementary Dataset 3.** Gene Ontology analysis performed in PANTHER for the targets identified for the different SATTs.

## TABLES

**Table T1.**
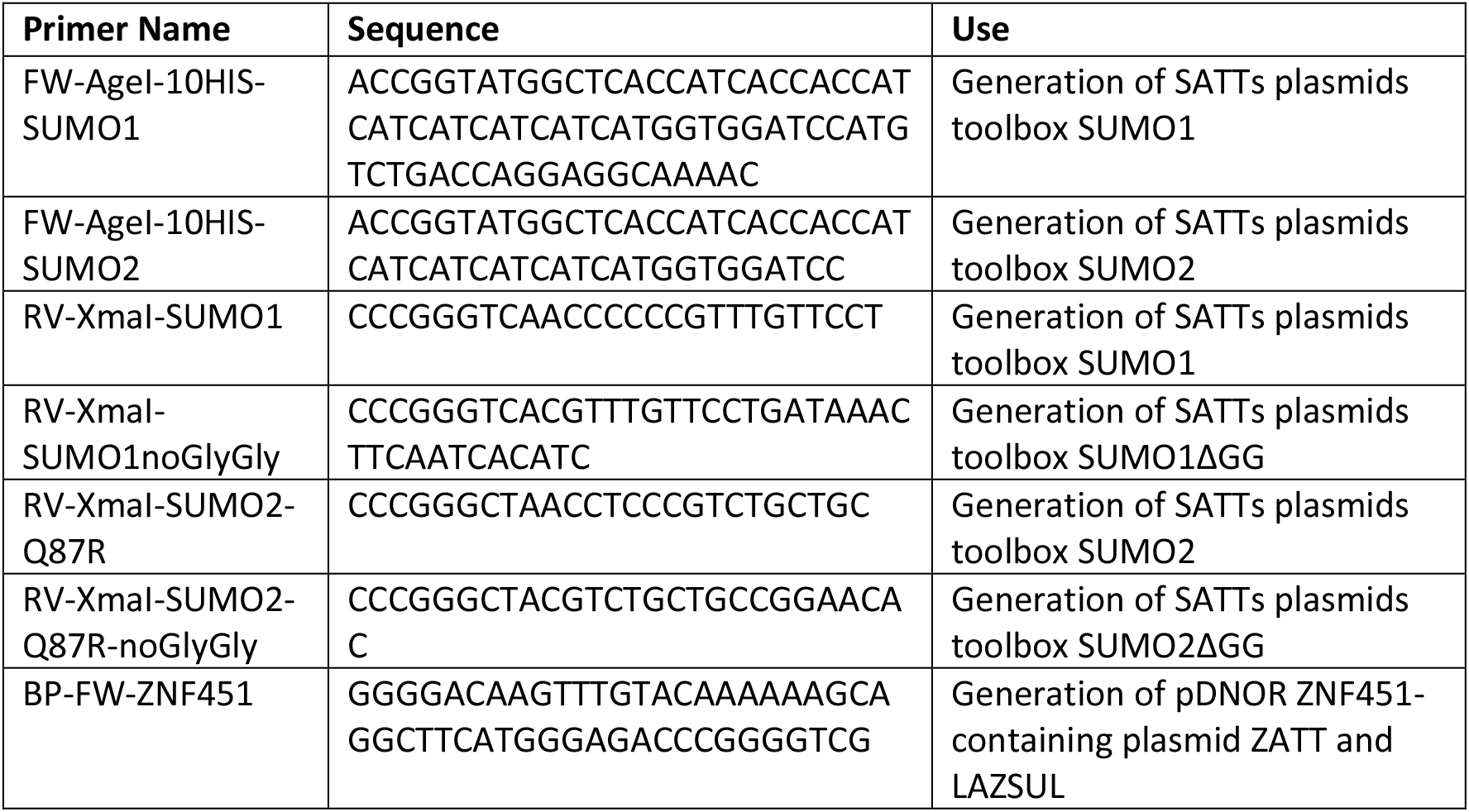
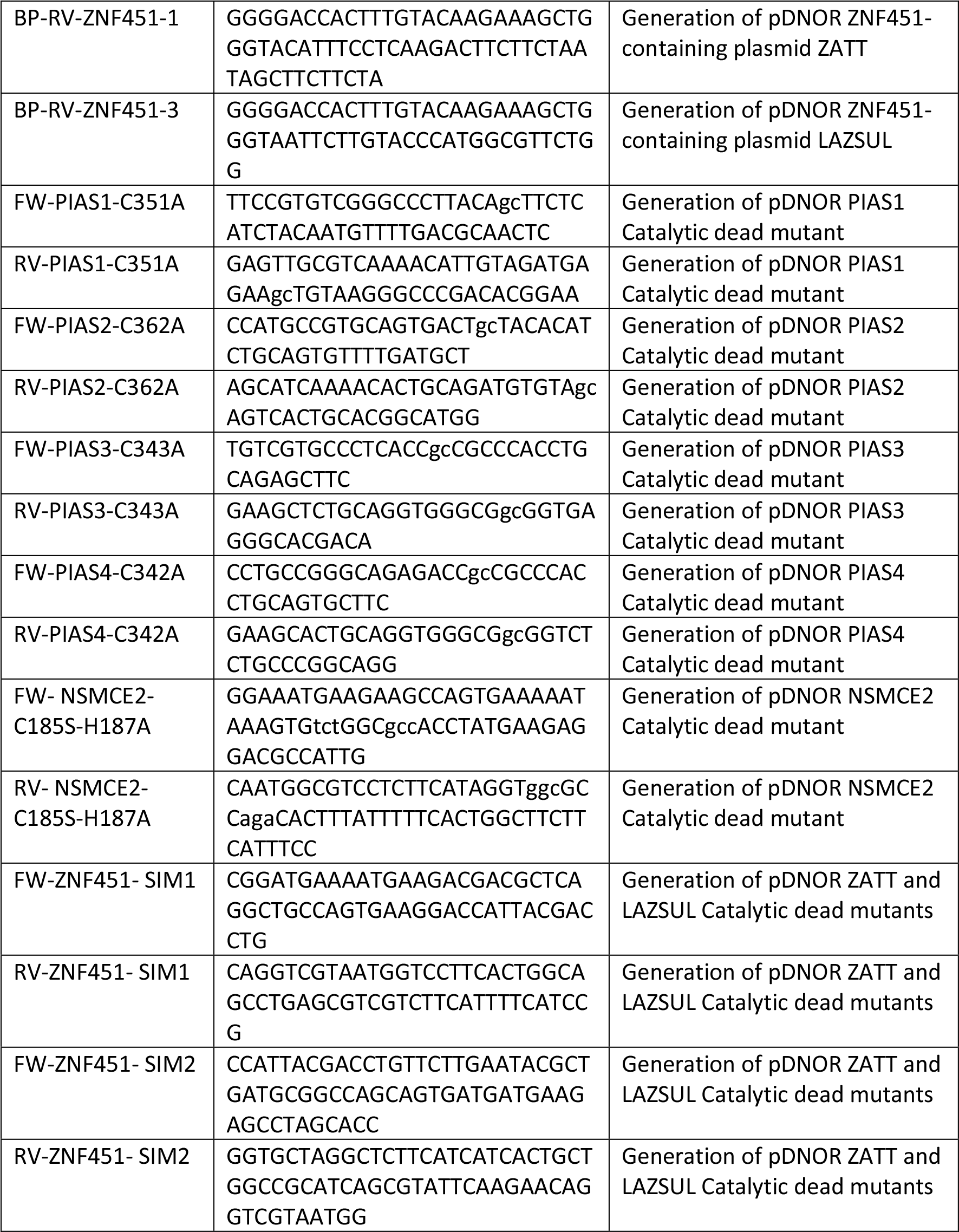
Primers

**Table T2.**
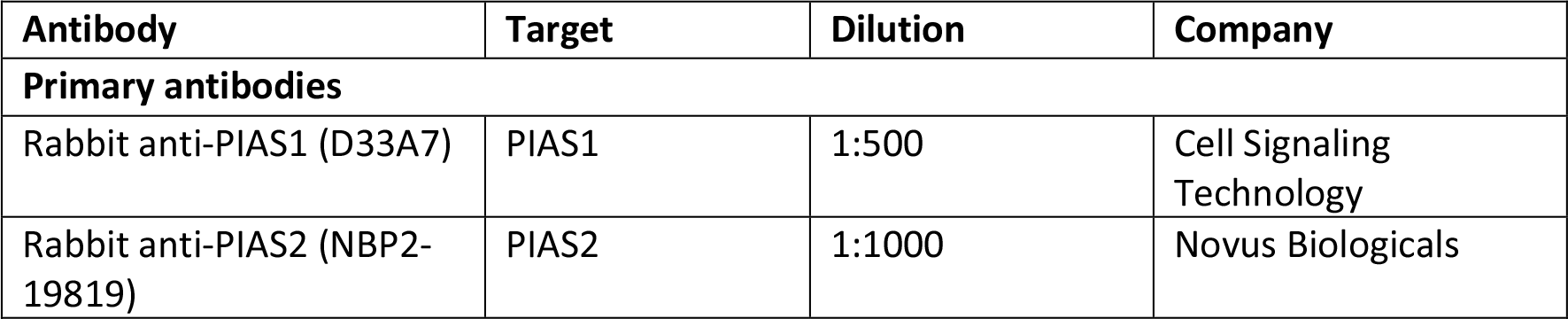
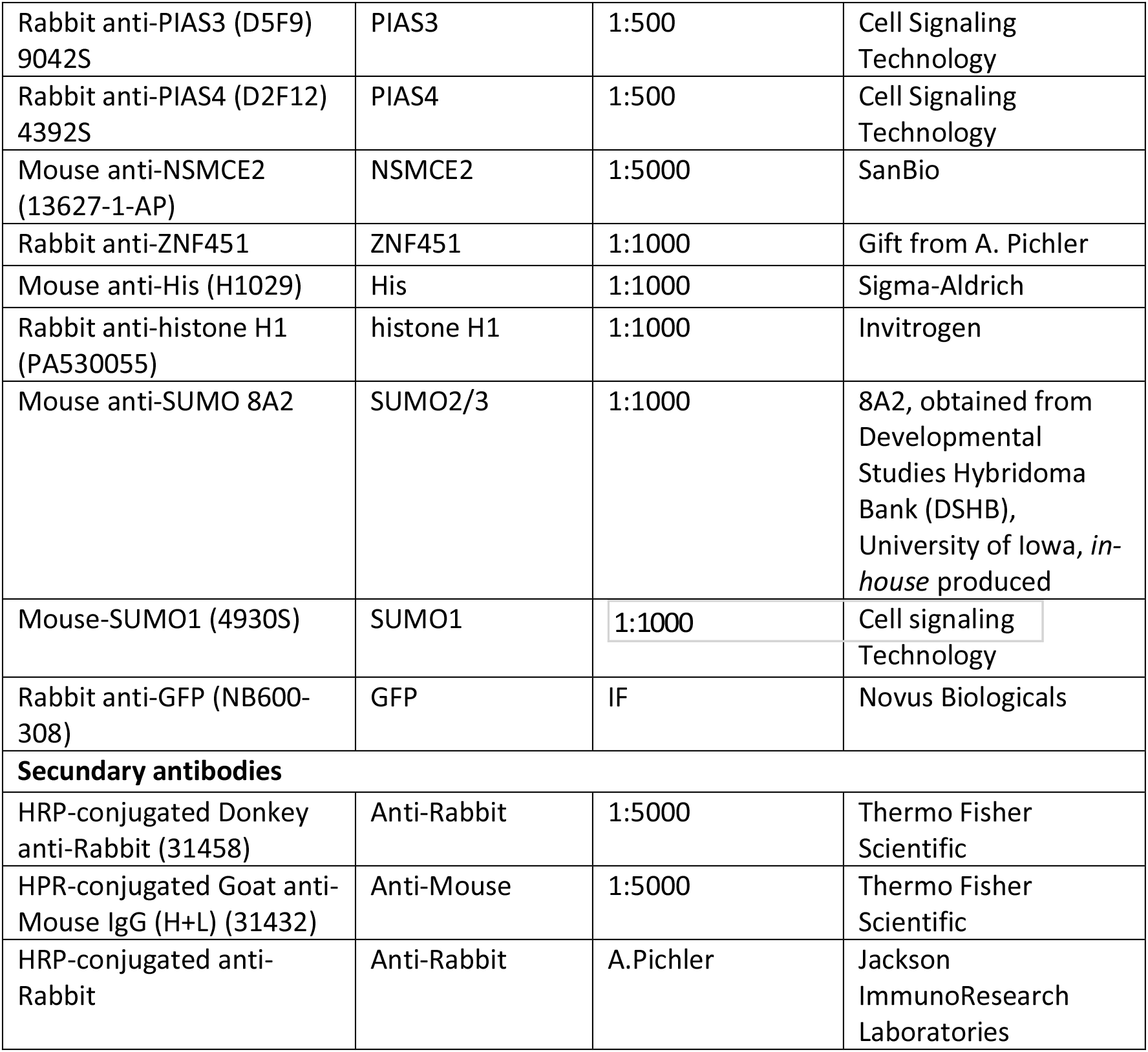
Antibodies

