## Supplementary Figures for "SUMO Activated Target Traps (SATTs) enable the identification of a comprehensive E3-specific SUMO proteome"

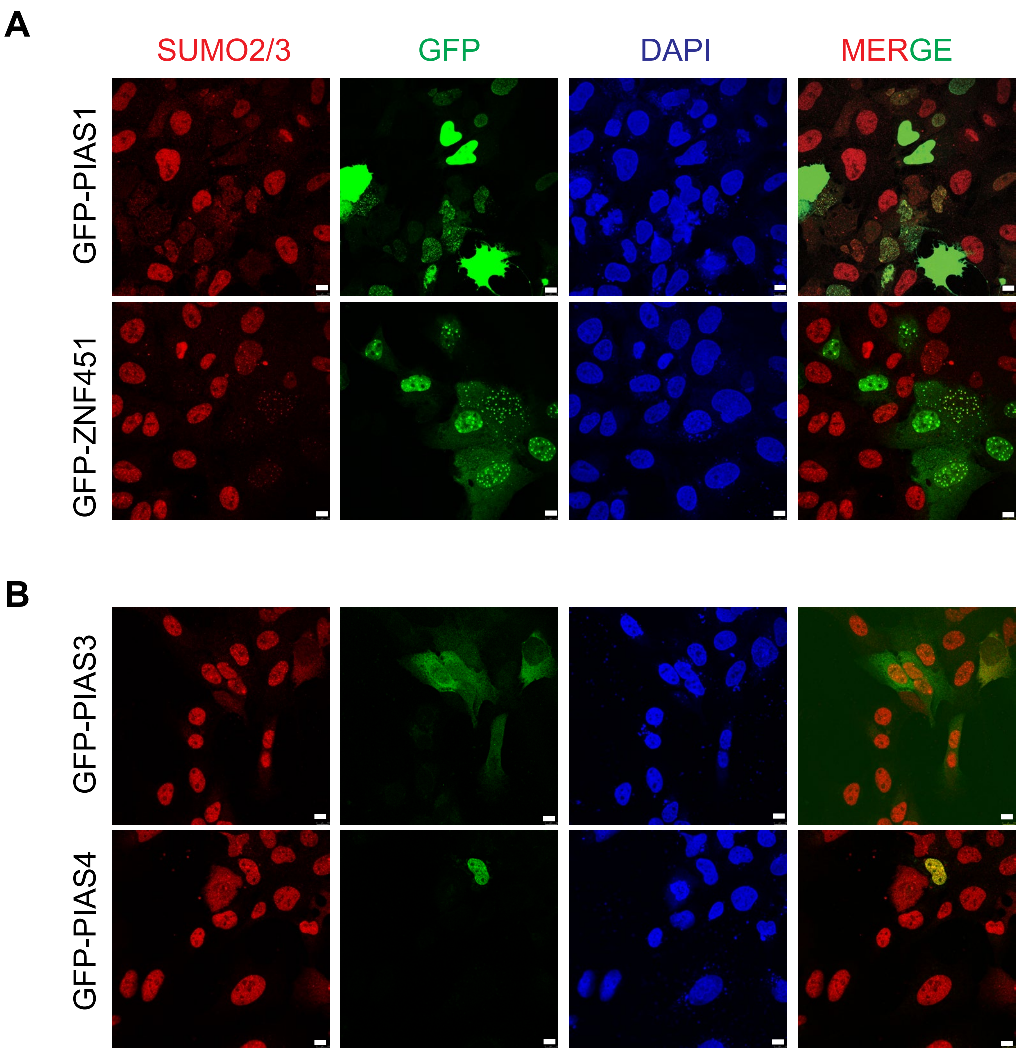


**Supplementary Figure 1.** Representative immunofluorescence images of U2OS cells transiently transfected with GFP-tagged E3s. Immunostained for SUMO2/3.

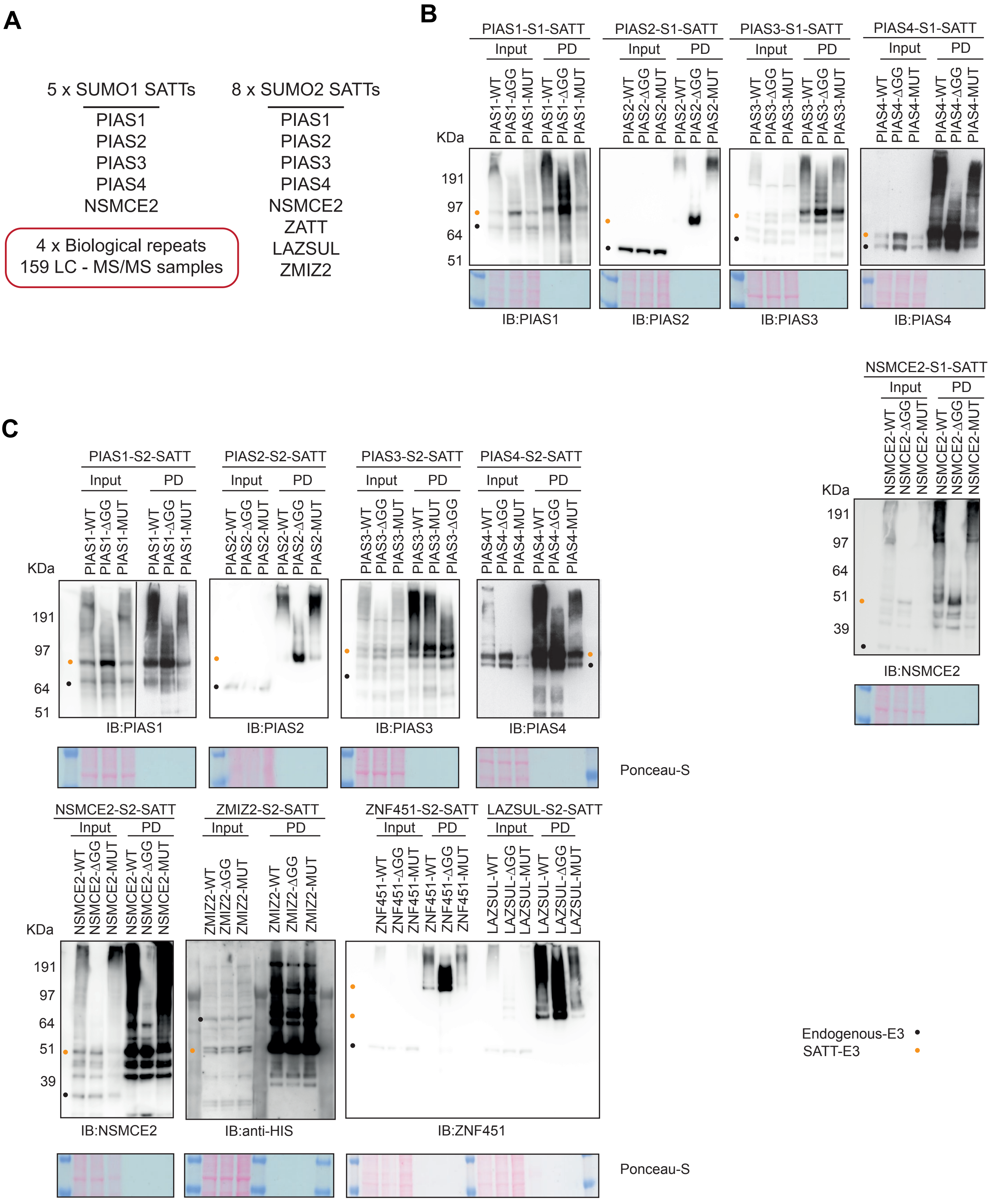


**Supplementary Figure 2. (A)** SUMO E3 SATTs investigated in this resource. **(B-C)** Immunobloting analysis of the SUMO1 (B) or $SUMO2_{Q87R}$ (C) SATTs samples. Antibodies used are indicated, Ponceau staining is included as loading control.
